# Intergenic regions contribute fitness advance by increasing recombination

**DOI:** 10.1101/2023.11.09.566346

**Authors:** Ryosuke Imai

## Abstract

Intergenic regions are common in eukaryotic genomes, yet they often appear to confer no direct fitness advantage. This study hypothesized that the primary function of these regions may be to enhance the frequency of meiotic recombination between genes, thereby reducing selective interference and improving the efficacy of selection. To test this hypothesis, computer simulations were conducted using the slim4 software to examine the impact of varying lengths of intergenic regions on total fitness. Populations with extended intergenic regions exhibited increased total fitness. Notably, the average fitness effects of both advantageous and deleterious mutations diminished, implying an enhanced selection efficiency, particularly for mutations with marginal fitness impacts. Total fitness effect of deleterious mutations increased despite the average fitness effect decreasing. Linkage between deleterious mutations could contribute to purging of slightly deleterious mutations.These results underscore the need for a more comprehensive examination of the relationship between recombination and intergenic regions to fully elucidate their evolutionary significance and contribution to genomic architecture.

**Article Summary:** Evolutionary significance of intergenic regions are still unclear. This study hypothesized that the primary function of these regions may be to enhance the frequency of meiotic recombination between genes, thereby reducing selective interference and improving the efficacy of selection. The simulation showed that populations with extended intergenic regions exhibited increased total fitness. The results poin t the way for future research about evolution of intergenic regions and the effect of genomic position of genes.

## Introduction

Intergenic regions, constituting a significant portion of the eukaryotic genome, largely consist of non-coding sequences such as transposable elements, which appear to offer no fitness advantage (Tenaillon *et al*. 2010; Ågren and Wright 2011). Paradoxically, despite their association with reduced growth rates and the potential for negative selection due to increased genome size (Hessen *et al*. 2010), these regions are ubiquitous across eukaryotes. The persistence of intergenic regions raises questions about their fitness implications and evolutionary significance.

This study proposes that intergenic regions may play a crucial role in enhancing meiotic recombination frequency between genes. By increasing the physical distance between genes, elongated intergenic regions could elevate recombination rates, thereby reducing gene linkage. Such reduction in linkage is theorized to alleviate background selection and enhance selection efficacy, as linkage can otherwise limit allelic combinations and impede natural selection (Peck 1994; Barton 1995, 2010; Otto 2021). When alleles are physically linked, their mutual influence can diminish the efficacy of selection. However, recombination serves as a mechanism to break these linkages, reducing selective interference in a stochastic process that restores selection efficiency under sexual reproduction (Nordborg *et al*. 1996; Keightley and Otto 2006). Therefore, it is postulated that the elongation of intergenic regions may mimic the effects of increased recombination rates, facilitating the reduction of selective interference through enhanced inter-gene recombination. As such, the length of intergenic regions could be a contributing factor to selection efficacy.

While recombination rates vary according to species, chromosomal positioning, and epigenetic states (Li *et al*. 2012; Peñuela *et al*. 2022), there is a general positive correlation between linkage map length and genome size, albeit with noted exceptions in non-flowering plant (Stapley *et al*. 2017; Brazier and Glémin 2022). Given that intergenic regions significantly contribute to genome size, they likely play a crucial role in shaping map lengths.

To test this hypothesis, computer simulations were conducted to assess the impact of intergenic region length on fitness enhancement. Comparisons were made between populations with varying intergenic region lengths using computer simulations, aiming to elucidate the importance of intergenic regions in evolutionary processes.

## Materials and methods

I conducted computer simulation using the software SLiM 4.0.1 (Haller and Messer 2022) to compare intergenic region length for fitness increase in different reproduction types. I assumed diploid hermaphrodite species with discrete non-overlapping generations and stable population sizes N = 10000. I compared different lengths of intergenic regions. The chromosome comprises 500 times repeat of the pair of 1000 bp coding regions and various lengths of intergenic regions among populations: 0, 100, 200, 300, 400, 500, 600, 700, 800 and 1000 bp.

I assumed that 99% of all mutations are deleterious and 1% are advantageous. Deleterious mutations have 0.2 dominance coefficient and gamma distribution fitness effect with mean of -1/2N, with a shape parameter (alpha) of 1.0. Advantageous mutations have 0.8 dominance coefficient and exponential distribution fitness effect with mean of 5/N. The mutation rate is 1.0*10e-7 and the recombination rate is 1.0*10e-8. I ran a simulation for 20N generations and measured the fitness effect of fixed mutations. I conducted 100 replicates on each condition.

## Result and Discussion

The simulation demonstrates a clear increase in total fitness concurrent with the elongation of intergenic regions. This enhancement is attributed to the increasing total fitness effects of fixed advantageous mutations. As the intergenic regions elongated, I observed a reduction in the average fitness effects of both advantageous and deleterious mutations. This suggests that selection efficacy for mutations with small fitness impacts is enhanced by longer intergenic regions. Previous study indicated that areas with high gene density may impede adaptive evolution through selective interference, unlike regions with longer intergenic stretches (Castellano et al. 2016). There is a difference, though, between within genomes and between populations, such findings corroborate the implications of my simulation. This study also implies that the length of intergenic regions may be instrumental in tuning selection efficiency via its effects on recombination within the genome level. A deeper investigation into the interplay between recombination and intergenic regions is warranted to fully understand how the evolution of genes and intergenic regions are interconnected.

Contrarily, the aggregate fitness effect of deleterious mutations has risen due to an expansion in the number of these mutations becoming fixed. This could be explained by two mechanisms in contexts of short intergenic regions: (1) strong deleterious mutations are purged alongside nearby alleles, irrespective of mutation type, and (2) clusters of slightly deleterious mutations, when closely linked, behave analogously to a single strong deleterious mutation, facilitating the elimination of mutations with minor fitness effects. The elongation of intergenic regions appears to mitigate these purging effects, leading to a higher fixation rate of slightly deleterious mutations. Should the number of advantageous mutations be limited, these dynamics might exert a more pronounced influence on overall fitness.

The simulation findings suggest that the length of intergenic regions influences the efficacy of selection and the pace of fitness advancement within a population, indicative of positive selection pressures acting upon intergenic region lengths. Nonetheless, the implications of intergenic region length evolution at the individual level remain unclear, primarily because these regions are generally neutral or nearly neutral, suggesting that natural selection may not directly target these intergenic region stretches. This is in line with Keightley and Otto (2006) (Keightley and Otto 2006), who posited that recombination rates might be subject to positive selection to circumvent selective interference. Changes in intergenic region length could have similar dynamics affecting modifier alleles as discussed in their study. Future research is essential to understand the role and intensity of natural selection on individual intergenic region lengths.

This study suggests that intergenic regions have indirect fitness advantages by increasing recombination. It could explain a part of evolutionary significance of intergenic regions and genome size variation.

## Acknowledgments

The author would like to thank to A. Satake and Y. Iwasa for insightful discussion.

## Conflicts of interest

None declared.

**Fig. 1.**
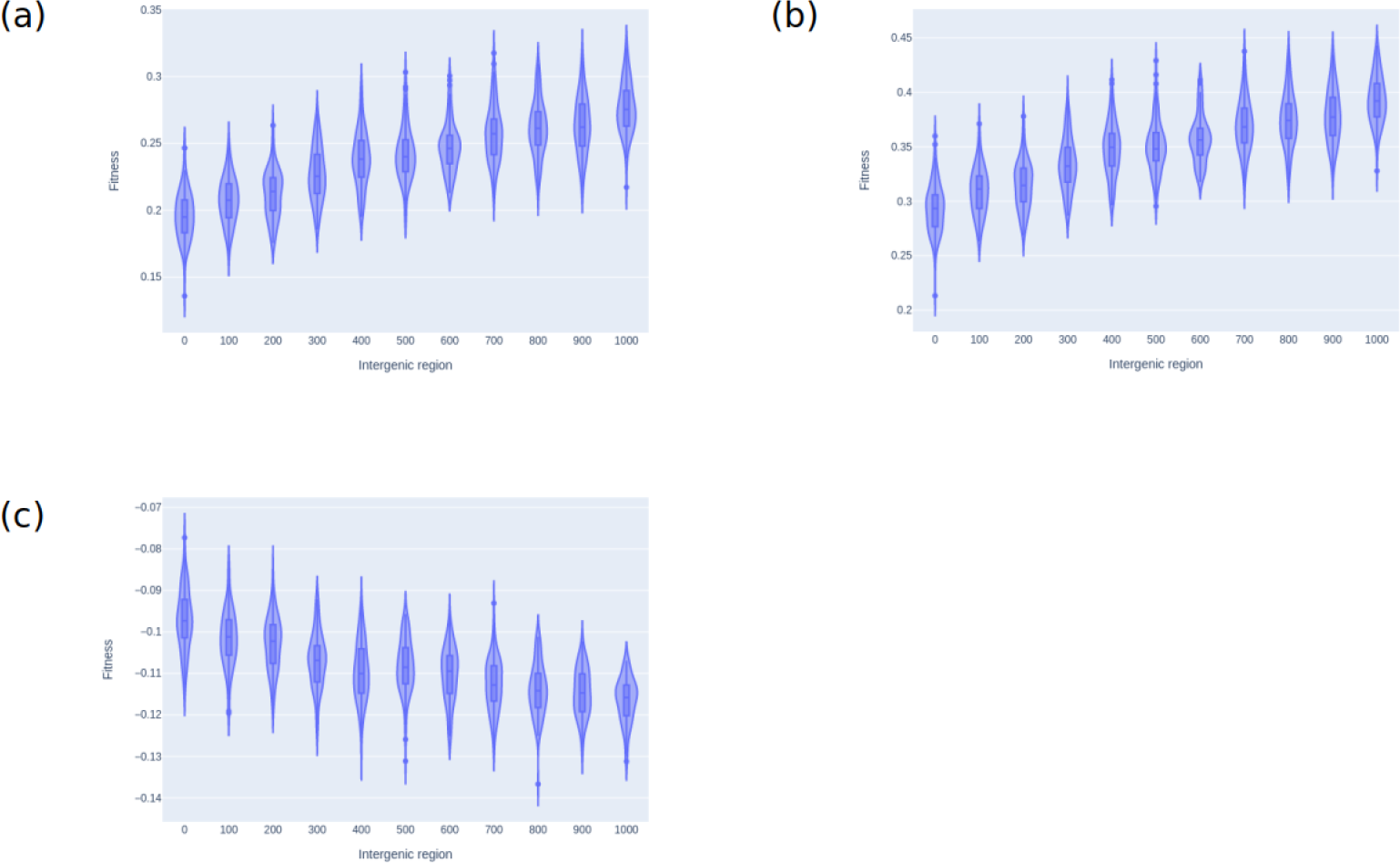
Total fitness effect of fixed mutations. Each violin plot and boxplot shows a total fitness effect of 100 times replication of each simulation. (a) Total fitness effect of all fixed mutations in each simulation. The Y axis is fitness and the X axis is intergenic region length per 100 bp coding region. (b) Total fitness effect of advantageous fixed mutations in each simulation. The Y axis is fitness and the X axis is intergenic region length per 100 bp coding region. (c) Total fitness effect of deleterious fixed mutations in each simulation. The Y axis is fitness and the X axis is intergenic region length per 100 bp coding region.

**Fig. 2.**
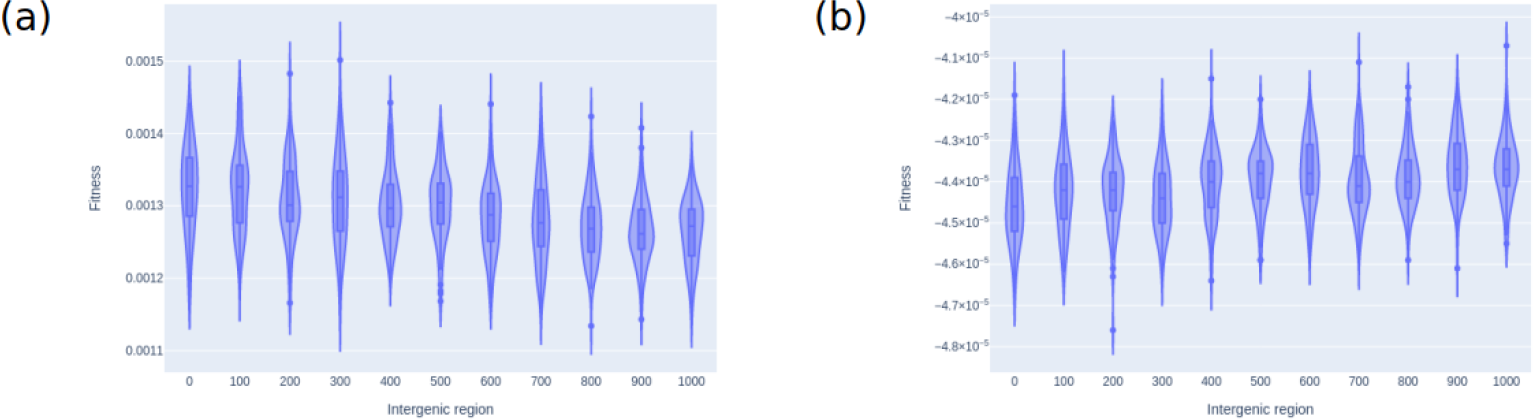
Average fitness effect of fixed mutations. Each violin plot and boxplot shows an average fitness effect of 100 times replication of each simulation. (a) Average fitness effect of advantageous fixed mutations in each simulation. The Y axis is fitness and the X axis is intergenic region length per 100 bp coding region. (b) Average fitness effect of deleterious fixed mutations in each simulation. The Y axis is fitness and the X axis is intergenic region length per 100 bp coding region.

**Fig. 3.**
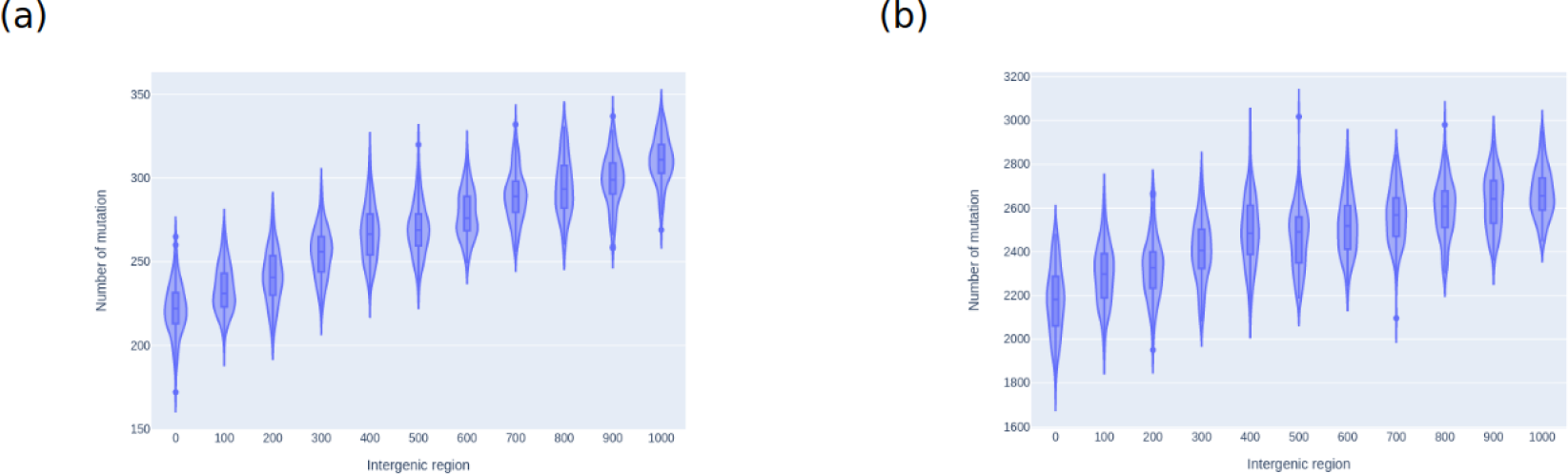
Number of fixed mutations. Each violin plot and boxplot shows a number of fixed mutations of 100 times replication of each simulation. (a) number of advantag eous fixed mutations in each simulation. The Y axis is number and the X axis is intergenic region length per 100 bp coding region. (b) number of deleterious fixed mutations in each simulation. The Y axis is number and the X axis is intergenic region length per 100 bp coding region.

## Reference

Ågren J. A., and S. I. Wright, 2011 Co-evolution between transposable elements and their hosts: a major factor in genome size evolution? Chromosome Res. 19: 777–786.

Barton N. H., 1995 Linkage and the limits to natural selection. Genetics 140: 821–841.

Barton N. H., 2010 Genetic linkage and natural selection. Philos. Trans. R. Soc. Lond. B Biol. Sci. 365: 2559–2569.

Brazier T., and S. Glémin, 2022 Diversity and determinants of recombination landscapes in flowering plants. PLoS Genet. 18: e1010141.

Haller B. C., and P. W. Messer, 2022 SLiM 4: Multispecies Eco–Evolutionary Modeling. Am. Nat. E000–E000.

Hessen D. O., P. D. Jeyasingh, M. Neiman, and L. J. Weider, 2010 Genome streamlining and the elemental costs of growth. Trends Ecol. Evol. 25: 75–80.

Keightley P. D., and S. P. Otto, 2006 Interference among deleterious mutations favours sex and recombination in finite populations. Nature 443: 89–92.

Li X., J. Zhu, F. Hu, S. Ge, M. Ye, et al., 2012 Single-base resolution maps of cultivated and wild rice methylomes and regulatory roles of DNA methylation in plant gene expression. BMC Genomics 13: 300.

Nordborg M., B. Charlesworth, and D. Charlesworth, 1996 The effect of recombination on background selection*. Genet. Res. 67: 159–174.

Otto S. P., 2021 Selective Interference and the Evolution of Sex. J. Hered. 112: 9–18.

Peck J. R., 1994 A ruby in the rubbish: beneficial mutations, deleterious mutations and the evolution of sex. Genetics 137: 597–606.

Peñuela M., J. J. Gallo-Franco, J. Finke, C. Rocha, A. Gkanogiannis, et al., 2022 Methylation in the CHH Context Allows to Predict Recombination in Rice. Int. J. Mol. Sci. 23. 10.3390/ijms232012505

Stapley J., P. G. D. Feulner, S. E. Johnston, A. W. Santure, and C. M. Smadja, 2017 Variation in recombination frequency and distribution across eukaryotes: pat terns and processes. Philos. Trans. R. Soc. Lond. B Biol. Sci. 372. 10.1098/rstb.2016.0455

Tenaillon M. I., J. D. Hollister, and B. S. Gaut, 2010 A triptych of the evolution of plant transposable elements. Trends Plant Sci. 15: 471–478.

